# Quantitative Analysis of Tissue Secretome Reveals the Diagnostic and Prognostic Value of Carbonic Anhydrase II in Hepatocellular Carcinoma

**DOI:** 10.1101/2020.01.15.905687

**Authors:** Xiaohua Xing, Hui Yuan, Hongzhi Liu, Xionghong Tan, Bixing Zhao, Yingchao Wang, Jiahe Ouyang, Minjie Lin, Aimin Huang, Xiaolong Liu

## Abstract

Early detection and intervention are key strategies to reduce mortality, increase long-term survival and improve the therapeutic effects of hepatocellular carcinoma (HCC) patients. Herein, the isobaric tag for relative and absolute quantitation (iTRAQ) quantitative proteomic strategy was used to study the secretome in conditioned media from HCC cancerous tissues, surrounding noncancerous and distal noncancerous tissues to identify diagnostic and prognostic biomarkers for HCC. In total, 22 and 49 secretory proteins were identified to be dysregulated in the cancerous and surrounding noncancerous tissues compared with the distal noncancerous tissues. Among these proteins, carbonic anhydrase II (CA2) was identified to be significantly upregulated in the secretome of cancerous tissues; correspondingly, the serum concentrations of CA2 were remarkably increased in HCC patients than that in normal populations. Interestingly, a significant increase of serum CA2 in recurrent HCC patients after radical resection was also confirmed compared with HCC patients without recurrence, and the serum level of CA2 could act as an independent prognostic factor for time to recurrence (TTR) and overall survival (OS). Regarding the mechanism, the secreted CA2 enhances the migration and invasion of HCC cells by activating the epithelial mesenchymal transition (EMT) pathway. Taken together, this study identified a novel biomarker for HCC diagnosis and prognosis and provides a valuable resource of the HCC secretome for investigating serological biomarkers.

## Introduction

Hepatocellular carcinoma (HCC) has a high incidence and mortality, making it the fifth most common malignant cancer worldwide [1]. Although surgical strategy was proven to be the most suitable option for HCC therapy, most HCC patients were not diagnosed or intervened until the advanced stage, rendering them unsuitable for surgical treatments [2] or having a poor prognosis after surgical excision. Currently, the five-year survival rate of HCC patients is less than 20%, while the five-year recurrence and metastasis rate is greater than 80% [3–6]. So far, alpha fetoprotein (AFP) and des-gamma-carboxy prothrombin (DCP) are the most widely accepted and clinically applied biomarkers for HCC diagnosis and monitoring. However, the sensitivity and specificity of AFP and DCP in the early diagnosis as well as prognosis evaluation of HCC are still insufficient [7–9]. Therefore, it’s urgently required to find novel biomarkers that are highly specific and sensitive to provide an early diagnosis and prognosis evaluation for HCC.

Proteomic-based technology has become a very useful and powerful analytical tool for biomarker screening [10–13]. A desirable biomarker for HCC diagnosis or monitoring should be able to be measured in body fluid samples such as serum and plasma [14], because these samples are low-cost, easy to collect and process, and are amenable to repeat sampling whenever it is necessary. Therefore, the serum and plasma are also the ideal targets for proteomic studies that aim to identify diagnostic or prognostic biomarkers for HCC [15,16]. However, the complex nature of serum and plasma, as well as their large dynamic concentration range of different proteins, significantly hinders the progress of proteomic-based biomarker screening.

Secreted proteins play important roles in signal transduction, cellular growth, proliferation, apoptosis and even in tumorigenesis, development, invasion and metastasis, and are ideal sources for biomarker screening [17]. Investigating the secretome of HCC tissues or cells may provide valuable information for relevant studies. Recently, the application of secretomics in screening diagnostic or prognostic protein biomarkers in HCC cell lines have been reported by many groups [18–20]. However, these results must still be clinically validated [21]. Therefore, it would be more straightforward and convincing to analyze the secretome of primary tumour tissue cultures to identify the diagnostic or prognostic biomarkers for HCC. For example, Yang et al. have established an *in vitro* tissue culture system for HCC and identified matrix metalloproteinase 1 (MMP1) as a diagnostic biomarker for HCC [22]; however, the influence of hepatitis B virus (HBV) infection was not analyzed in this study.

In the present study, we collected serum-free culture media (CM) from the tissue cultures of tumour tissues, surrounding non-tumoral tissues and distal non-tumoral tissues of HCC patients and analyzed the secretome to identify potential diagnostic and prognostic biomarkers for HCC via an iTRAQ-based quantitative proteomic approach. Meanwhile, the sensitivity, specificity and clinical significance of the identified biomarkers was also carefully validated in a large-scale HCC patient cohort by ELISA assay and targeted proteomics of parallel reaction monitoring (PRM). Furthermore, the corresponding molecular mechanisms of the identified biomarker was also carefully explored.

## Results and Discussion

### Cells in the *in vitro*-cultured tissues were alive and secretory

To ensure that the cells in the *in vitro*-cultured tissues were alive and that the secretome was not contaminated by intracellular proteins, a series of analyses were performed (Figure 1A). We used haematoxylin and eosin (HE) staining to evaluate the cell morphology changes of tissues cultured for 0 day, 1 day and 2 days. As revealed in Figure 1B, the HE-stained tissue sections showed the corresponding characteristic anatomical details of HCC cancerous, surrounding noncancerous and distal noncancerous tissues in all of the respective cultures. With the extension of incubation time, the cells in the cultured tissues were starved and showed necrosis or apoptosis due to the lack of nutrients. The morphology of tissues cultured for 1 day was still very similar to that of the fresh tissues (cultured 0 days). By contrast, the number of cell nuclei in tissues cultured for 2 days significantly decreased due to cell necrosis or apoptosis during the culture process (Figure 1B). TdT-mediated dUTP nick-end labeling (TUNEL) staining was further used to evaluate the apoptosis rates of tissues cultured for 0, 1 and 2 days. As revealed in Figure 1C, the tissues cultured for 2 days had a significantly higher apoptosis rate compared with tissues cultured for 0 or 1 days. In addition, the proteins extracted from culture supernatants with different culture times were examined by SDS-PAGE. As shown in Figure 1D, the molecular mass distribution of the extracted proteins was significantly changed with the increase of culture time. Furthermore, western blot assays clearly demonstrated the prevention of contamination by intracellular proteins (Figure 1E). Taken together, these results suggested that the 1 day culture time was the optimal time point for collecting culture supernatants.

**Figure 1.**
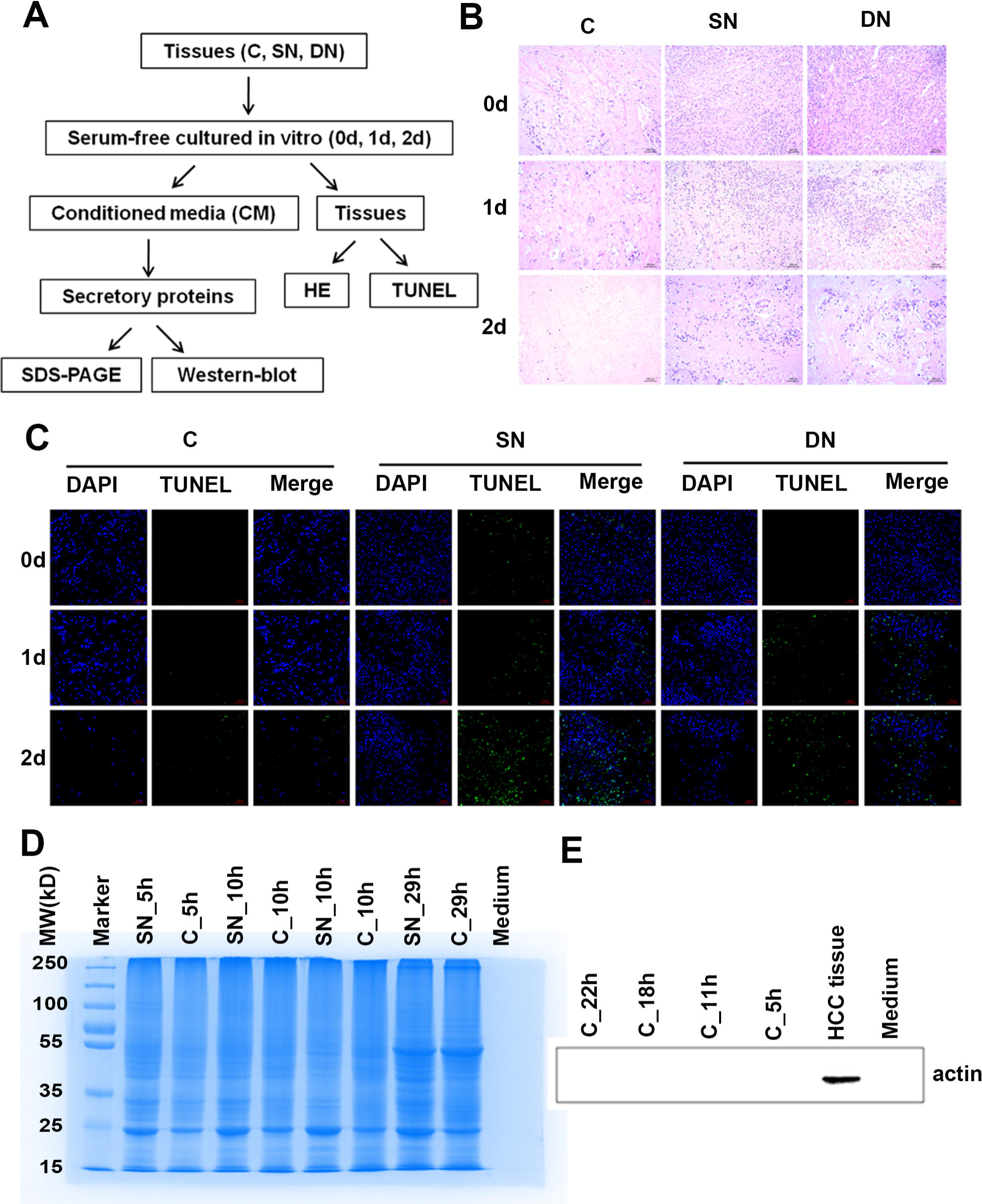
Strict quality control of the tissue secretome. **A.** The quality control workflow for the cultured tissues and the proteins secreted proteins in the supernatant. **B.** HE staining of cultured tissues. The three columns were the staining of cancerous HCC tissues (left column, 200 ×), surrounding noncancerous tissues (middle column, 200 ×) and distal noncancerous tissues (middle column, 200×), cultured for 0, 1 and 2 days, respectively. **C.** TUNEL staining of cultured tissues. We examined the densities of DAPI (blue) and FITC (green) for every tissues. **D.** The molecular weight distribution of secretory proteins through SDS-PAGE. **E.** Western blot of secretory proteins.

### The secretome characteristics were comprehensive and reasonable

iTRAQ labeling combined with mass spectrometry was applied to investigate the entirety of secretome changes in primary HCC tissues from patients. The features of the HCC patients in the current study were listed in Table S1. Total proteins extracted from the supernatant collected from the culture medium of HCC tissues and their surrounding and distal noncancerous tissues were analyzed using 2D LC-MS/MS, as shown in Figure 2. We quantified 2388 proteins in total using Scaffold_4.3.2, of which 1312 proteins were annotated or predicted as secretory proteins, accounting for 54.9% of the total quantified proteins. This result covered 75.7% of the secretory proteins previously reported by Yang et. Al [19] (Figure S1A), and the percentage of secretory proteins was much higher than in the human protein database (23%) (Figure S1B). Among these proteins, 936 secretory proteins were overlapped in the 5 different replicates, which accounted for 71.4% of the quantified secretory proteins. The complete list of identified secretory proteins was shown in Table S2. The detailed features of identified secretory proteins, including the NM-score, MetazSecKB characteristics, isoelectric point (pI), molecular weight (MW), hydrophobicity and quantification results were also included in the list. The MS-based proteomic data were deposited in the integrated proteome resources of iProX public repository (iProX Accession: IPX0001425001) at http://www.iprox.org/page/MSV022.html [23].

**Figure 2.**
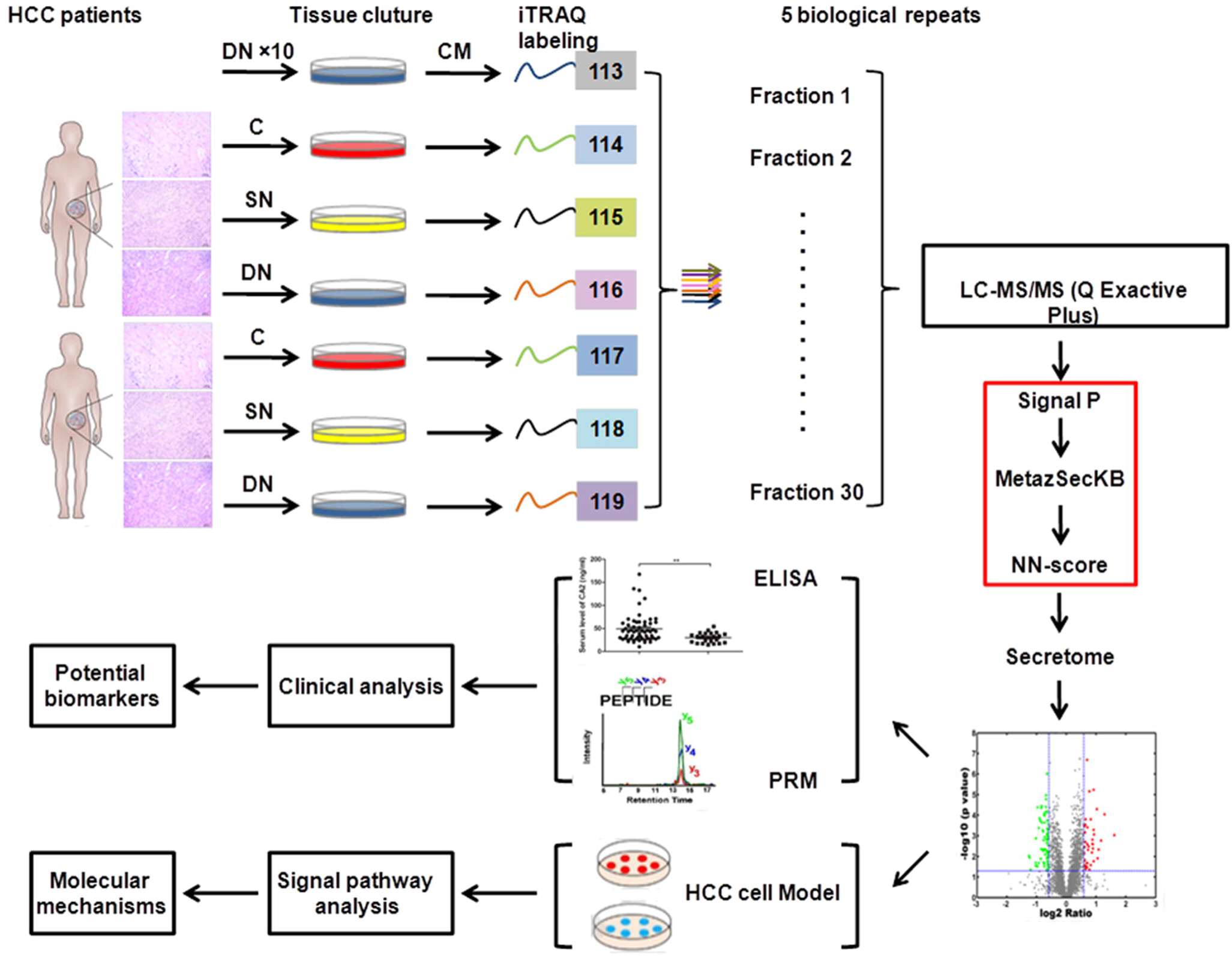
Experimental workflow. The HCC tissues, surrounding noncancerous tissues and distal noncancerous tissues were cultured *in vitro*, and the conditioned medium was collected to extract secretory proteins. Secretory proteins were digested with trypsin, directly labeled using iTRAQ-8plex and analyzed through 2D LC-MS/MS. The target proteins screened by bioinformatics were then verified *in vitro* and *in vivo* to find potential biomarkers of HCC and to investigate the molecular mechanisms of HCC recurrence.

The MW distribution of the secretory proteins ranged from 7396 to 628685 Da, with a primary range of 10∼40 kD, indicating smaller molecular weight for the secretory proteins (Figure 3A). The pI values ranged from 3.67 to 12.56 and were mainly in the range of 4.4∼7.6, which coincides with the microenvironment of liver tissues (Figure 3B). The hydrophobicity of the proteins ranged from 1.1 to 3.3 and was mainly in the range of 1.6∼2.0, which implied the enrichment of membrane or transmembrane proteins (Figure S1C).

**Figure 3.**
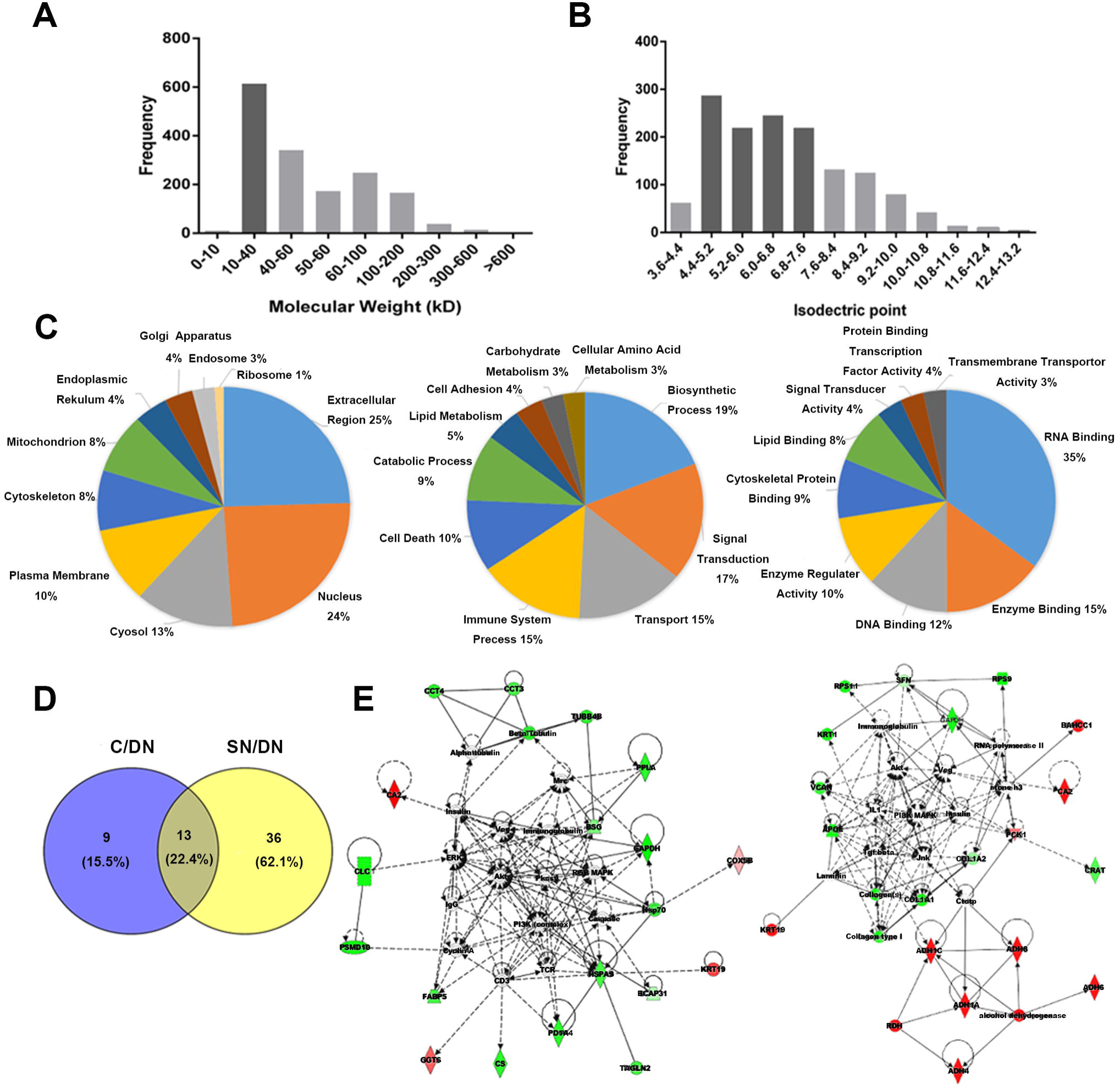
Objective and reasonable features of the HCC tissue secretome. The distribution of (**A**) molecular weight and (**B**) isoelectric point of the identified secretory proteins. GO analysis of the (**C**) cell component, (**D**) biological processes and (**E**) molecular functions of the secretory proteins. (**F**) Venn diagrams of secretory proteins for the two comparisons (C/DN group and SN/DN group). The signaling pathway networks involved in the (**G**) C/DN group and (**H**) in SN/DN group.

We also investigated the GO annotations containing cell components, molecular functions and biological processes of the secretory proteins. Cell component category showed that the secretory proteins were primarily extracellular, which suggests the excellent purity of HCC the tissue secretome (Figure 3C). The biological process category indicated that these secretory proteins were mainly involved in biosynthetic process, signal transduction, and transport process (Figure 3D). The molecular function category indicated that these secretory proteins played major roles in RNA binding, enzyme binding, DNA binding and transmembrane transporter activities (Figure 3E). These results were consistent with thoes of previously reported secretory proteins from HCC tissues [19].

### Various functions of secretory proteins in different HCC-related tissues

For proteomic analysis, HCC tissues, surrounding noncancerous tissues and distal noncancerous tissues were obtained from 10 patients who underwent surgery, and the samples were divided into 3 groups as follows: cancerous tissue group (C group); surrounding noncancerous tissue group (SN group) and distal noncancerous tissue group (DN group). In the current study, the differentially abundant proteins had the same change tendency in all 10 biological replicates and presented a fold change of approximately ± 1.5 (log_2_ 0.58) in at least 5 biological replicates with a *p* value < 0.05. Under this standard, there were 22 differentially abundant proteins in C/DN group (Table S3), and 49 differentially abundant proteins in SN/DN group (Table S4). The numbers of overlapped differentially abundant proteins among two comparisons was showed by the Venn diagram in Figure 3F. Among these differentially abundant proteins, 13 differentially abundant proteins were shared in two groups. As the GO annotation analysis indicated, these proteins mainly participated in extracellular matrix organization, extracellular structure organization, and tissue morphogenesis (Figure S1D), which might be a contributing factors for HCC development. According to the result, 9 dysregulated proteins were found in C/DN group but not in SN/DN group. These proteins were the mainly participated in the disorder of primary metabolism (Figure S1E), which is tightly linked to the development of HCC, and even invasion, infiltration and metastasis. Meanwhile, 36 dysregulated proteins were found in SN/DN group but not in C/DN group. Similarly, we found these proteins were the mainly participated in acute-phase response, acute inflammatory response and post-transcriptional regulation on gene expression (Figure S1F), suggesting that the surrounding noncancerous tissues are distinctly different from the distal noncancerous tissues. Interestingly, these processes were all related to primary metabolism, suggesting that changes in primary materials might play a crucial role in the occurrence of HCC.

To further study the potential molecular mechanisms of the occurrence and development of HCC, IPA analysis was applied to investigate the signaling pathways in which the dysregulated proteins participated. The results showed that the two comparisons (C/DN group and SN/DN group) indeed had specific signaling pathways, although they also had common shared signaling pathways. As the IPA investigation showed, the dysregulated proteins in C/DN group were mostly involved in ERK/MAPK signaling, while the dysregulated proteins are mainly participated in PI3K-Akt signaling pathway in SN/DN group. There are 11 proteins participated in ERK/MAPK signaling (2 up-regulated and 9 down-regulated) (Figure 3G). The ERK/MAPK pathway can transduce extracellular signals through intracellular signal transduction cascades to finally control the expression of proteins that regulate tumorigenesis and aggressive behaviours [24,25]. The dysregulation of ERK/MAPK signaling pathway in C/DN group revealed that the secreted factors from HCC cancerous tissue might modulate the tumour micro-environment to exert important roles in tumorigenesis.

In SN/DN group, 20 proteins (4 up-regulated and 16 down-regulated) participated in PI3K-Akt signaling (Figure 3H). PI3K-Akt signaling plays a crucial role in the regulation of inflammation and metabolism [26–28]. Hence, alterations of the PI3K-Akt pathway might be closely linked to the occurrence and development of tumours. The dysregulation of PI3K-Akt signaling pathway in the secretory environment of HCC adjacent noncancerous tissues compared with that of distal noncancerous tissues suggested the importance of changing microenvironment in tumorigenesis. Targeting these important effectors in tumour microenvironment might be a promising therapeutic strategy.

### CA2 might be a valuable biomarker for HCC diagnosis

According to the IPA analysis, we found that carbonic anhydrase II (CA2) was dysregulated between the two comparison (C/DN group and SN/DN group). CA2, a zinc metal enzyme, carries out the reversible hydration of carbon dioxide, and plays a key role in adjusting the pH of tumour microenvironment [29–34]. It is frequently abnormally expressed in different cancers [35–41]. Here, the CA2 relative intensity of the reporter ions of the 8-plex iTRAQ reagent in the MS/MS spectra were in good agreement with the protein levels. As revealed in Figure S2A, the levels of CA2 were remarkably increased in C/DN group and SN/DN group (114>115>116 and 117>118>119). Therefore, CA2 might be a potential interesting biomarker for HCC diagnosis and prognosis prediction. Here, we further verified the clinical significance of CA2 on the diagnosis and prognosis of HCC in two additional serum cohorts via the PRM targeted proteomic method and traditional ELISA assays.

The CA2 serum concentration was monitored in 49 HCC patients and 23 healthy volunteers through PRM. There were 9 identified peptides from CA2 in total, 7 of which were overlapped in the five 8-plex iTRAQ experiments, and 1 of which had a mis-cleavage site. Therefore, only 6 peptides from CA2 could be used for the PRM experiment, and the annotated spectra and detailed PSM information for these 6 identified CA2 peptides were provided in Table S5 and Figure S3. For the skyline results, the peak contributions of the individual fragment ions from the unique peptide was showed in Figure S2B, and the representative quantification information based on peak areas of the peptides including endogenic and synthetic heavy peptides was displayed in Figure S2C. The CA2 serum concentration in HCC patients was remarkably higher than that in normal populations (*p* < 0.01, 7.01 and 13.72 pg/mL average serum CA2 in healthy volunteers and HCC patients, respectively). The receiving operating character (ROC) curve analysis of CA2 revealed that the area under the curve (AUC) was 0.715 for HCC patients relative to the healthy volunteers (Figure 4A). These results were consistent with the data from the proteomic studies, indicating that CA2 might be a valuable biomarker for HCC diagnosis.

**Figure 4.**
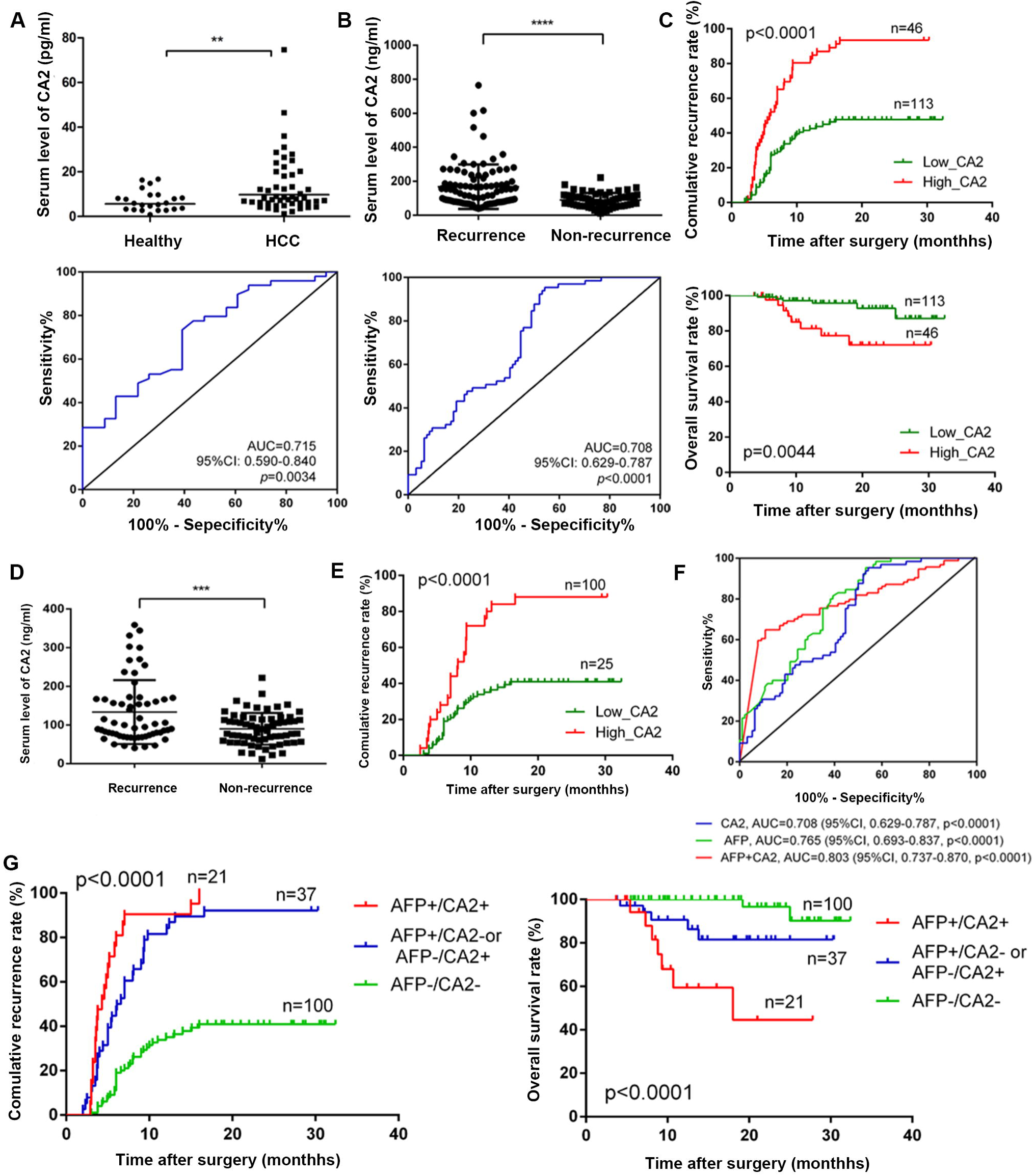
Clinical values of serum CA2 in HCC diagnosis and prognosis. **A.** Distribution and ROC curve of serum CA2 levels in HCC patients and healthy volunteers from the training cohort through PRM. **B.** Distribution and ROC curve of serum CA2 levels in HCC patients, with and without recurrence, from the validation cohort assessed by ELISA. **C.** TTR and OS were compared between the high- and low-CA2 groups in the validation cohort by Kaplan-Meier analysis. **D.** Distribution of serum CA2 levels in AFP-negative HCC patients. **E.** TTR was compared between the high- and low-CA2 groups in AFP-negative HCC patients by Kaplan-Meier analysis. **F.** AUC of the CA2/AFP combination in HCC patients. **G.** Prognostic values of the CA2/AFP combination in HCC patients.

### CA2 is a novel prognostic biomarker for HCC

According to our previous study, the serum levels of CA2 were steadily increased from 3 months to 9 months after radical resection in the patients with short-term recurrence compared with the patients without recurrence (Figure S4A). According to these data, we expect to find an appropriate time point after radical resection to predict the recurrence of HCC. The serum samples taken 5 months after surgery were collected blindly. An ELISA assay was used to analyze the association between the CA2 serum concentrations and the prognosis for 159 HCC patients, containing 94 relapsed patients and 65 relapse-free patients. As revealed in Figure 4B, serum CA2 was remarkably upregulated in HCC patients with recurrence compared with HCC patients without recurrence (*p* < 0.01). According to the ROC curve, the AUC reached 0.708 in the validation cohort, indicating that CA2 might be a HCC prognostic biomarker. To evaluate the prognostic significance of serum CA2 in HCC, we further studied the overall survival and recurrence rate of patients with low and high serum levels of CA2 using the 159 samples. The median concentration (105.3 ng/mL) was used as the optimal cutoff value to stratify patients into low (≤ 105.3 ng/mL) and high (> 105.3 ng/mL) CA2 groups. As shown by the Kaplan-Meier analysis, HCC patients with a higher serum CA2 level had a higher recurrence rate than those with a low serum CA2 level. Furthermore, the overall survival (OS) of HCC patients who had a high serum CA2 level was remarkably shorter than those with a low CA2 level (Figure 4C). Furthermore, we repeated this study using the training cohort of 49 HCC patients and obtained identical results (Figure S4B). In addition, we found a strong negative linear correlation between serum CA2 levels and the recurrence time, indicating that the higher the serum CA2, the shorter the recurrence time (Figure S4C). These results suggested that the serum CA2 level might be a novel prognostic biomarker for HCC.

### CA2 might be a novel prognostic biomarker for AFP-negative HCC patients

As the gold standard for clinical diagnosis and monitoring in HCC, AFP was remarkably higher in relapsed HCC patients than that in patients without relapse (*p* < 0.001) (Figure S4D). Although the recurrence rate was substantially lower while the overall survival was remarkably higher in AFP-negative patients than that in AFP-positive patients in general (AFP-negative and AFP-positive patients were divided according to clinical detected AFP serum concentration, 20 ng/mL) (Figure S4E), some patients with negative serum AFP still experienced rapid recurrence or metastasis (Figure S4F). Therefore, there is an urgent need to identify novel biomarkers that can predict the recurrence/metastasis in AFP-negative HCC patients. Interestingly, among the AFP-negative HCC patients, the serum concentration of CA2 in relapsed patients was remarkably higher than that in relapse-free patients (*p* < 0.001) (Figure 4D). Furthermore, in the AFP-negative HCC patients, the recurrence rates were also higher in patients with high CA2, suggesting the prognostic values of CA2 in AFP-negative patients (Figure 4E). The overall survival rate did not show significant differences between the high-CA2 and low-CA2 groups (Figure S4G).

### Combination of serum CA2 and AFP improves prognostic performance in HCC

According to the above results, we confirmed that CA2 had a prognostic value in AFP-negative patients. We then jointly considered the CA2 and AFP serum levels. As shown in Figure 4F, when CA2 and serum AFP were considered together, the AUC reached to 0.803 for the combination of CA2/AFP (AUC of 0.708 for CA2 and 0.765 for AFP), suggesting a clinical values for the combination of these 2 markers. Next, we further evaluated the prognostic value of CA2/AFP in combination for HCC patients in the validation cohort. As shown in Figure 4G, the cumulative recurrence rate in patients in the low-CA2 and AFP-negative group was remarkably lower than that of patients in the high-CA2 and AFP-positive group. Correspondingly, the overall survival of patients in the low-CA2 and AFP-negative group was substantially higher than that of patients in high-CA2 and AFP-positive group. CA2-positive and AFP-negative or CA2-negative and AFP-positive patients had a median OS and TTR in the validation cohorts. Taken together, these results suggested an improved prognostic value when using the serum CA2/AFP levels in combination for HCC patients.

### Serum CA2 was significantly correlated with tumour number and BCLC stage

To further study molecular mechanisms of CA2 on the prognosis of HCC, we further checked the association between serum CA2 and various clinicopathologic features of HCC patients. As revealed in Table S6, the Pearson’s chi-square test has indicated that higher serum CA2 level was significantly correlated with more tumour number (*p* = 0.026) and more advanced BCLC stage (*p* = 0.042), but did not associated with other clinicopathologic features. These results suggested that CA2 might be a metastasis/recurrence-related protein in HCC.

### Secreted CA2 increases the migration and invasion abilities of HCC cells by activating of EMT signaling

To investigate the molecular mechanism of extracellular CA2 in HCC recurrence/metastasis, exogenous recombinant CA2 protein (obtained from Abcam) was used to examine its influences on the migration and invasion ability of HCC cells using a trans-well strategy. As shown in Figure 5A, exogenous recombinant CA2 treatment of MHCC97L cells significantly promoted the migration and invasion of cells compared to control cells with the same concentration of BSA treatment (*p* < 0.05), suggesting that extracellular CA2 enhances cell migration and invasion in HCC, which could explain the observed clinical data. The epithelial-mesenchymal transition (EMT) is well known to be responsible for tumour metastasis. As shown in Figure 5B, the MHCC97L cells treated with exogenous recombinant CA2 exhibited a spindle-like fibroblastic morphology, while the corresponding control cells were round with a more epithelial morphology. Such morphological changes indicated that the secreted CA2 might be involved in the EMT process. To further investigate the involved molecular mechanisms, the related key markers of EMT were also analyzed. As shown in Figure 5C, the addition of exogenous recombinant CA2 upregulated N-cadherin (a mesenchymal marker) and downregulated E-cadherin (a epithelial maker); meanwhile, the expression levels of vimentin and Zeb1 (EMT-promoting transcription factors) were also significantly upregulated by the addition of exogenous recombinant CA2. Taken together, these results demonstrated that the secreted CA2 promoted the EMT transition to further modulate the migration and invasion of HCC cells, in turn affecting HCC metastasis.

**Figure 5.**
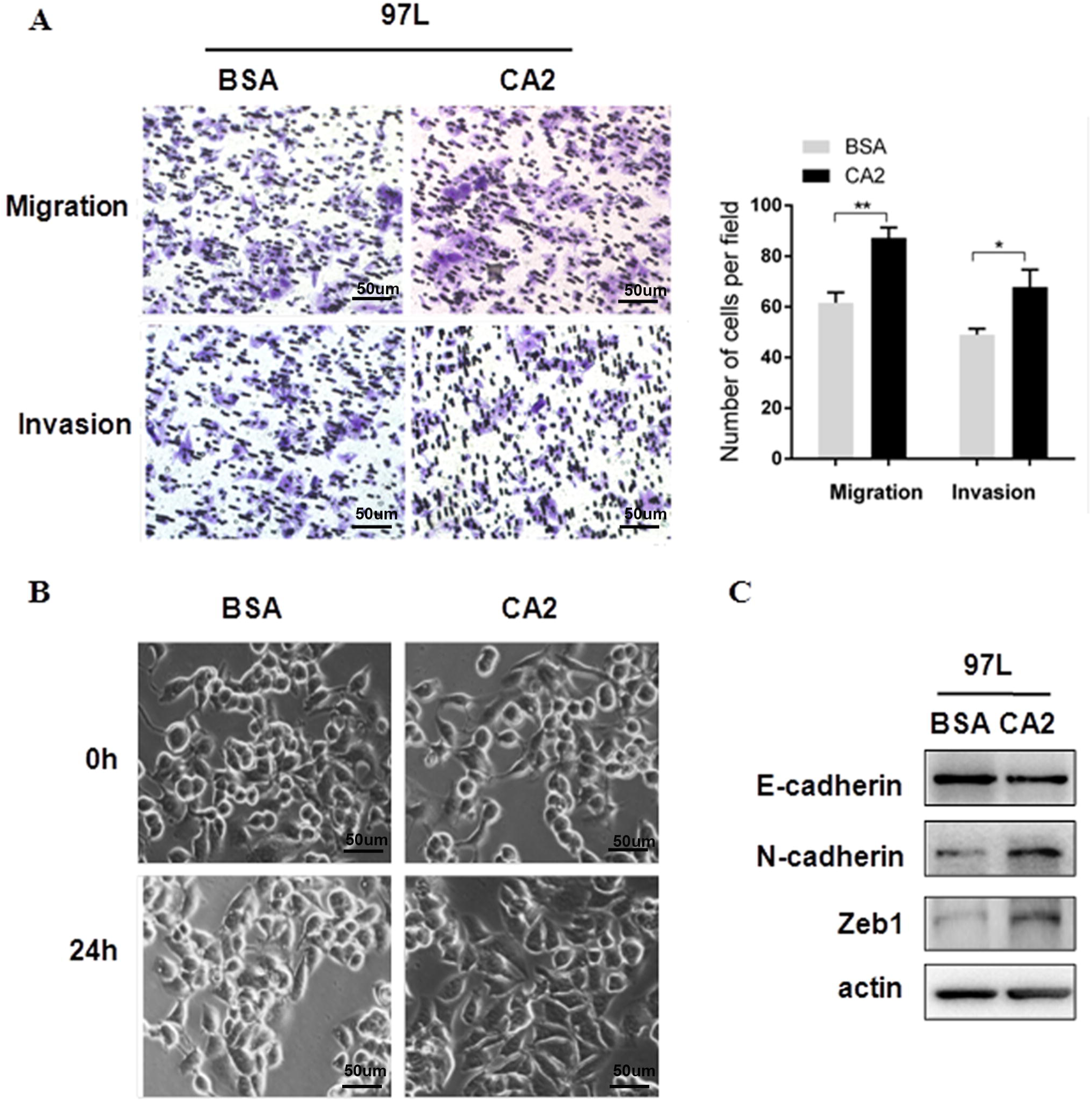
Extracellular CA2 promotes cell migration and invasion through activating EMT. **A.** Representative images and quantification results of cell migration and invasion in MHCC97L cells after adding exogenous recombinant CA2. **B.** Cellular morphology of HCC cells with the addition of exogenous recombinant CA2; Magnification, ×200; Scale bar, 50 μm. **C.** Western blot analysis showed that adding exogenous recombinant CA2 downregulated the expression of E-cadherin, while upregulating the expression of Zeb1 and N-cadherin.

### Intracellular CA2 might perform opposite functions from its extracellular form

We also studied the intracellular levels of CA2 in HCC patients. In this experiment, 28 paired HCC tissues and their corresponding noncancerous tissues were detected by western blot. As revealed in Figure 6A and B, the intracellular CA2 expression was remarkably decreased in HCC tissues when compared with their paired noncancerous tissues, and the representative images of CA2-immunostained HCC tissues were shown in Figure S5. Meanwhile, the downregulation of intracellular CA2 in tumour tissues was further confirmed by immunostaining using in-house TMAs containing 75 HCC tissues and their paired noncancerous tissues (Figure 6C). The results were exactly opposite from those of the extracellular CA2 in the serum of HCC patients, suggesting that intracellular and extracellular CA2 might carry out opposite functions [42]. However, the involved molecular mechanisms should be further explored.

**Figure 6.**
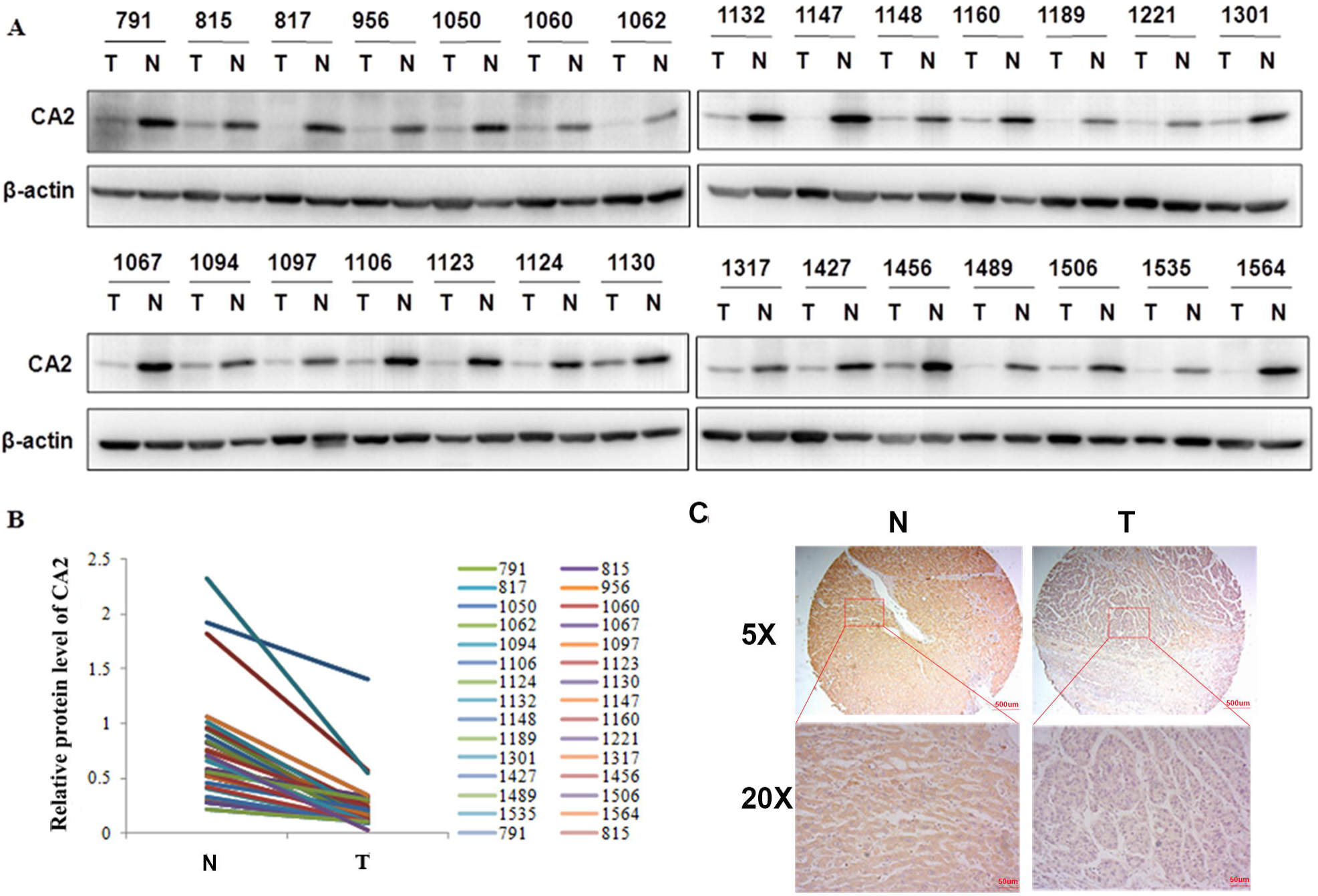
Expression of intracellular CA2 was opposite of the extracellular form. **A.** Intracellular CA2 expression in HCC tissues and their paired surrounding non-tumoral tissues. **B.** Intracellular CA2 expression was deceased in HCC tissues compared with their paired surrounding non-tumoral tissues. **C.** IHC analysis of the intracellular CA2 expression in HCC tissues and their paired surrounding non-tumoral tissues on in-house TMAs.

## Conclusion

Herein, we applied the iTRAQ-based quantitative proteomic approach to investigate the secretome of the primary culture of HCC tissues, and identified secreted CA2 as a diagnostic and prognostic biomarker for HCC. In particular, CA2 showed good predicative performance in AFP-negative HCC patients, and the combination of CA2 and AFP improved the sensitivity and specificity of HCC prognosis. Regarding the mechanism, extracellular CA2 might regulate the HCC cell migration and invasion by targeting the EMT signaling pathway to affect HCC patients’ prognosis. This secretome investigation enabled us to identify a novel HCC diagnostic and prognostic biomarker. The information from this study constitutes a valuable resources for further HCC investigation and for identifying potential serological biomarkers.

## Materials and methods

### Patients and follow-up

In total, 293 HCC patients and 23 healthy volunteers were enrolled in the current study. All of the HCC patients underwent surgical procedures at the Mengchao Hepatobiliary Hospital of Fujian Medical University (Fuzhou, China). Computer tomography (CT) scanning, angiography and ultrasonography (US) were used to monitor the absence of intrahepatic recurrence and metastasis in the residual liver. All patients met the enrolment eligibility criteria as follows: (1) The patients were diagnosed with HCC through pathological examination after operation; (2) Serum hepatitis B surface antigen (HBs Ag) was positive and hepatitis B surface antibody (HBs Ab) and hepatitis C virus (HCV) were negative before surgery; (3) Standard radical resection was performed: no distal metastasis was found before or during the operation; intraoperative ultrasound examination revealed no other liver lesions; no obvious tumour thrombus was found in the hepatic portal vein or primary venous branch; postoperative pathological examination showed no cancer cells at the cutting edge; no recurrence/metastasis was found in the ultrasound and CT examination 2 months after the operation; (4) Serum AFP was increased before operation and decreased to normal level two months after surgical operation; (5) The patient had not received any other intervention or treatment before surgery.

For the proteomic analysis, HCC tissues, surrounding noncancerous tissues and distal noncancerous tissues were obtained from 10 patients who underwent surgical operation. Serum from 49 HCC patients before hepatectomy and 23 healthy volunteers in a health screening program at the Mengchao Hepatobiliary Hospital were collected for PRM analysis. Another validation serum cohort from 159 HCC patients after hepatectomy with long-term follow-up that ended with death was used for the ELISA assay. The collection of serum samples was strictly controlled following the in-house standard operating procedure, which was established on the basis of previous studies [43]. In addition, a total of 75 formalin-fixed and paraffin-embedded HCC tissues and their adjacent noncancerous tissues from patients who underwent surgical resection were collected to fabricate the tissue microarrays (TMAs) for immunohistochemistry investigation.

This project was approved by the Institution Review Board of Mengchao Hepatobiliary Hospital of Fujian Medical University. Informed consent was obtained from each participant before the operation. The use of clinical specimens was completely in compliance with the “Declaration of Helsinki”.

### Tissue culture and quality control *in vitro*

Following surgical operation, the primary tissue specimens were immediately transferred into PBS on ice and sent to the laboratory within 30 min. The tissues were then rinsed and cut into 2-3 mm^3^ pieces, and extensively washed several times at room temperature with PBS to eliminate major blood and serum contaminants. Subsequently, the samples were transferred to 10 cm cell culture dishes and incubated in serum-free DMEM medium supplemented with 1% penicillin (cat No. 15070-063, Gibco, USA) and streptomycin (cat No. 15070-063, Gibco, USA) at 37 °C. For protein extraction, the supernatants were centrifuged at 100 g for 2 min at 4 °C to further remove all remaining cells and debris; the samples were then concentrated with a 3 K cutoff centrifugal filter device (cat No. UFC500396, Millipore, USA) and stored at -80 °C until further analysis.

Culture medium was collected every 4 hours in following 72 h period (a total of 18 time points). The proteins extracted from every time point were subjected to 10% SDS-PAGE to analyze the molecular mass distribution and protein blotting was performed to determine any contamination by intracellular proteins. At the same time, the cultured tissues at every time point were evaluated by histological observations. The cultured tissues were fixed in 10 % neutral formaldehyde, and paraffin sections were made in the conventionally way. One slide was stained with HE (cat No. D006, Nanjing Jiancheng Bioengineering Institute, China) and observed under an optical microscope (AE2000, Motic, USA); another slide was biotin-labeled and stained by TUNEL to detect cell apoptosis during the culture period. Finally, the slides were observed under a confocal microscope (LSM 780, Carl Zeiss, Germany).

### Bottom-up proteomics and data analysis

The proteomic studies and data analyses were modified from our previously reported protocols [44]. Briefly, the thiol groups of the above extracted protein samples from the culture supernatant of 3 groups were alkylated with 50 mM iodoacetamide (cat No. I6125, Sigma, USA) (30 min in dark at room temperature) after reduction with 8 mM DTT (cat No. D0632, Sigma-Aldrich, USA) (55 °C, 1 h). Then, the proteins were then precipitated by ice-cold acetone (5 times volume), and re-dissolved in 100 mM tetraethyl-ammonium bromide (TEAB) (cat No. 90360, Sigma, USA). Subsequently, 100 μg of each proteins sample was digested by trypsin (cat No. V511, Promega, USA) using filter aided sample preparation (FASP), and the peptides were labeled with the 8-plex iTRAQ reagent (cat No. 4381663, AB SCIEX, USA) as follows: C group, SN group and DN group were labeled with 114, 115 and 116, respectively; and one biological repetition of the above 3 groups was labeled with 117, 118 and 119, respectively. A, B, C, D and E were defined as the 5 independent iTRAQ 8-plex labeling repetitions. In addition, the DN group samples mixed in equal amounts were labelled with 113 and included in every 8-plex labelling reaction as an internal standard to balance each 8-plex labeling. The labeled peptides were mixed in equal amounts in every 8-plex labeling and were desalted by Sep-Pak Vac C18 cartridges (cat No. WAT023590, Waters Corporation, USA). The samples were then dried by the vacuum centrifuge (cat No. 7310038, LABCONCO, USA) for further use.

The peptide mixture was separated using an offline LC system (Acquity UPLC, Waters Corporation, USA) via high-pH separation. High-pH (pH = 10) separation was performed in a reverse-phase column (C18, 1.7 µm, 2.1×50 mm) (cat No. 186002350, Waters Corporation, USA) using a 20 min linear gradient from 5% B to 35% B (A: 20 mM ammonium formate (cat No. 70221, Sigma, USA) in water, B: 20 mM ammonium formate added in 90% ACN (cat No. A998, Thermo Fisher, USA), ammonium hydroxide (cat No. 17093, Sigma, USA) was used to adjust the pH). Finally, a total of 30 fractions were collected and 2 equal-interval fractions were combined to reduce the MS running time: for example, 1 and 16, 2 and 17, etc. [45]. In total,15 fractions were dried and subsequently separated on a Nano-LC system (Nano-Aquity UPLC, Waters Corporation, USA) with a 75 min linear gradient from 2% D to 40% D (C: 0.1% formic acid in water, D: 0.1% formic acid in ACN), which was performed on an analytical column (C18, 75 μm*15 cm, 3 μm) (cat No. 164534, Thermo Fisher Scientific, Germany). Next, the peptides were detected by mass spectrometry (Q-Exactive, Thermo Fisher Scientific, Germany) with 2.1 kV electrospray voltage at the mass spectrometer inlet. In addition, 70 K mass resolution was applied in the full-scan of MS spectra processing (m/z 350-1200), and 17.5 K resolution was used in the following 15 sequential MS/MS scans of high energy collisional dissociation (HCD). In all studies, 1 microscan was recorded by using a dynamic exclusion of 30 s.

The data processing strategy was also carried out according to our previous publications, with some modification [44]. The LC-MS/MS data acquisition was processed using Proteome Discoverer (version 1.4, Thermo Fisher Scientific, Germany) and searched using the Sequest HT (version 2.5.1, Matrix Science, United Kingdom) search algorithms against the human database (Uniprot, 20,264 entries, released at April 10, 2014). Proteome Discoverer were searched with trypsin for protease digestion and maximally we only allowed 2 missed cleavages. 10 parts per million (ppm) parent ion tolerance and 0.02 Da fragment ion mass tolerance were set according to MS precision. Fixed modifications included the iTRAQ modification of lysine residues and the peptide N-terminus, as well as carbamidomethylation of cysteine, while the variable modifications included iTRAQ labeling of tyrosine and oxidation of methionine. The peptide and protein identification false discovery rate (FDR) was calculated using the percolator algorithm against a decoy database and was set at 1% FDR. For protein quantitation, we only considered peptides that were unique to a certain given protein. The fold change between different samples was calculated by the ratio between iTRAQ reporter ion intensity and MS/MS spectra (m/z 113-119). The ratios were derived from criteria as follows: 20 ppm fragment ion tolerance was applied for the most confident centroid peak. In Sequest HT, the quantitative protein ratios were normalized and weighted by the median ratio. In addition, Scaffold (version 4.3.2, Proteome Software Inc., USA) was applied to verify the MS/MS-based peptides and identified proteins. The Scaffold was used to analyze and evaluate the quantitative results of iTRAQ, as well as to combine multiple quantitative experiments according to the designed internal standard in each 8-plex iTRAQ experiment for subsequent analysis. In detail, the raw data was searched by Proteome Discoverer using Sequest HT search algorithms, and the generated file was imported into the Scaffold software. The integrated information of the five 8-plex iTRAQ experiments could then be obtained directly by grouping. In the same way, a peptide FDR < 1% and a protein probability > 99.0% were accepted.

Two complementary methods were combined to further characterize the secretory proteins. First, MetazSecKB, a secretome proteome knowledgebase of metazoan, was performed to screen secretory proteins, which is the most direct source by which to characterize the secretory proteins. Second, SecretomeP (version 2.0, DTU Health Tech, Denmark), a sequence-based prediction strategy for mammalian secretory proteins, was used to classify the secretory proteins. The classical secreted proteins could be correctly predicted with an N-terminal signal peptide; non-classical secretory proteins without an N-terminal signal peptide could be correctly predicted as secretory according to an NN-score > 0.6. Those proteins without an N-terminal signal peptide and that also had a low NN-score were not identified as secretory proteins.

The GO annotation and signaling pathway investigation of differentially abundant proteins were performed using the free online tool DAVID (http://david.abcc.ncifcrf.gov/). The enriched signaling pathways of the dysregulated proteins were analyzed using Ingenuity Pathways Analysis (IPA) (version 7.5, Ingenuity Systems, Inc., USA).

### Targeted proteomics and data analysis

Before applying the targeted proteomic method of parallel reaction monitoring (PRM) analysis, the serum samples were immunoaffinity depleted of the 14 most abundant proteins using IgY14 LC20 (cat No. 5188-6557, Agilent, USA). The depletion was performed on an Agilent 1260 HPLC (1260 Infinity, Agilent, USA) system following the manufacturer’s protocol. The depleted serum was concentrated and the buffer was exchanged to 100 mM TEAB using Amicon 3 K concentrators (cat No. UFC500396, Millipore, USA). The procedures for serum protein denaturation, reduction, alkylation and digestion were described above.

For PRM analysis, the unique peptides for CA2 were synthesized by Fmoc solid-phase synthesis with isotope-labeled on the carboxyl side of the amino acid lysine (^13^C_6_,^15^N_2_) and purified by HPLC with purity > 99% (Anhui Guoping Pharmaceutical Co., Ltd, China). The heavy-labeled peptides were mixed and spiked into the tryptic digests of serum proteins at a concentration of 30 fg/μL. Detailed information for the unique CA2 peptides were displayed in Table S5. The unique peptide for CA2 should preferably have a narrow, symmetrical chromatographic peak, be 8-25 amino acids in length, ionize efficiently, provide a stable and intense signal without any modification, and not elute at the beginning or end time.

The PRM analyses were also performed on a mass spectrometer (Q Exactive Plus, Thermo Fisher Scientific, Germany). LC separation was executed with a trap column (C18, 75 μm*2 cm, 3 μm) (cat No. 164946, Thermo Fisher Scientific, Germany) and an analytical C18 column (C18, 75 μm*15 cm, 3 μm) (cat No. 164534, Thermo Fisher Scientific, Germany) on a nano-LC system (EASY-nLC 1000, Thermo Fisher Scientific, Germany). 2 μL of the tryptic digests and depleted serum samples were injected, and a gradient of 2% D up to 35% D over 30 min was applied. In all experiments, a PRM scan was performed at m/z 200, with a resolution of 70 K, an AGC target of 1 × 10^6^, a maximum injection time of 200 ms and an isolation window of ± 2. In addition, a normalized collision energy of 27% was used for ion dissociation, and a fixed first mass at 120 m/z was set. The inclusion list including the m/z and corresponding retention times of precursor peptides of interest were displayed in Table S5.

The PRM raw data was analyzed with Skyline software (version 4.2, University of Washington, USA), and a 5-minute window was used in the Skyline software. The sum of peak areas of the 5 most intense product ions was considered for protein quantification. Some ions needed to be excluded: for example, those ions that did not match the retention time of other monitored ions, or that showed interference signals, or that gave intense signals at other retention time.

### ELISA assay

The serum levels of CA2 in HCC patients were analyzed using an ELISA assay kit (cat No. LS-F29508, LifeSpan BioScience Inc., USA) following the manufacturer’s instructions. Briefly, the standard proteins and patient serum samples were diluted with the sample dilution buffer and then 50μL of the diluted standards proteins and samples were added to 96-well plates. Next, 100 μL of HRP-conjugate solution was carefully added to each well and incubated for 1 hour at 37 °C. Then, the plate was then carefully washed 4 times with PBS, and 100 μL of chromogen solution (1:1 solution A and solution B) was added to each well and further incubated for 15 min at 37 °C. Finally, 50μL of stop solution was added to stop the reaction, and the optical density at 450 nm was measured by a spectrophotometer (M5e, Molecular Devices, USA).

### Western blot

Western blotting was performed according to a previous publication [44]. Briefly, proteins were separated on a 12% SDS-PAGE, and the gel was transferred to a nitrocellulose membrane (cat No. HATF00010, Millipore, USA). The membrane was then blocked in 5% BSA (cat No. A7906, Sigma-Aldrich, USA) at room temperature for 2 hours, then further incubated in primary antibody against CA2 (1/1000 dilution, cat No. ab226987, Abcam, USA) at 4°C overnight. The corresponding secondary antibody was incubated with the membrane at RT for 1 hour after which the blots were carefully washed 4 times with TBST buffer and revealed using enhanced chemiluminescence reagents (cat No. 34080, Thermo Scientific, USA) and visualized by autoradiography.

### Immunohistochemistry (IHC)

Immunohistochemistry (IHC) was carried out on HCC TMAs according to a previous publication [46]. Briefly, after pre-treating at pH6 and blocking with peroxidase, the sections were further incubated with primary antibody against CA2 (1/50 dilution, cat No. ab226987, abcam, USA) for 0.5 hour. The sections were then treated with envision FLEX/HRP reagent for 20 min and then washed and stained by the envision FLEX-DAB chromogen (cat No. DM827, Dako, Denmark) and by Mayer’s Hematoxylin (Lille’s Modification) Histological Staining Reagents (cat No. ab220365, Abcam, USA) for 3 minutes. They were then washed in distilled water for 5 minutes. All pathological sections underwent double-blind scoring as follows: negative (0), weak (1), strong (2) or very strong (3) by two different pathologists double-blindly. A score of 0-1 indicates low expression, and a score of 2-3 indicates high expression.

### Cell migration and invasion assays

The cell migration and invasion assays were carried out according to our previously published protocols [44]. The cell migration ability was investigated by using transwell units with 8 μm pores (cat No. 3428, Corning Costar, USA), and cell invasion was investigated by using 24-well transwell inserts that had been precoated with matrigel with 8 μm pores (cat No. 354480, BD Biosciences, USA). A total of 1×10^5^ cells from the indicated treatments were cultured in serum-free medium in the upper chamber. To induce cell invasion and migration, DMEM supplemented with 10% FBS was placed in the lower chamber. The cells that adhered to the lower surface were then fixed with paraformaldehyde after 18 h of incubation and then stained by crystal violet (0.1%). The cells that adhered to the bottom surface were carefully counted within 5 different views under a microscope using 20 × magnification.

### Statistical analysis

A threshold for the iTRAQ ratio was set to screen secretory proteins whose abundance was substantially changed in the tumour tissue group compared with its corresponding noncancerous tissue group. The proteins were defined as differentially dysregulated if the iTRAQ ratio was higher than 1.5 or lower than 0.67 in at least 5 patients, and they also must had the same direction of alteration in all 10 biological replicates. The iTRAQ ratio was based on a comparison of the reporter ion intensities in the tumour tissue group compared to the corresponding noncancerous tissue group.

SPSS 19.0 was used for statistical analysis. Two-tailed paired Student’s t-test was used to compare quantitative data between two groups. Fisher’s exact test was applied to analyze the relationships between CA2 and clinical-pathological features. The Kaplan-Meier method was applied to calculate survival curves, while the differences were determined by using a log-rank test. In all analyses, *p* < 0.05 was recognized as statistically significant.

## Authors’ contributions

XHX, AMH and XLL designed the project, analyzed the data and revised the manuscript; XHX executed the study, performed the tissue culture, bottom-up proteomics and targeted proteomics assays, statistical analysis, and drafted the manuscript; HY and XHT carried out the IHC and transwell assays; HZL carried out the ELISA assay and participated in the statistical analysis; BXZ and YCW participated in the study; JHO performed the western blotting; MJL was involved in the clinical sample collection. All authors have carefully read and approved the submitted manuscript.

## Acknowledgements

This work was supported by the National Natural Science Foundation of China (Grant No. 81702910 and Grant No. 81672376); the Educational Commission of Fujian Province (Grant No. 2018B013); and the Natural Science Foundation of Fujian Province (Grant No. 2017J01159 and Grant No. 2016J01417).

## Supplementary material

**Figure S1 Features of the dysregulated secretory proteins**

**A.** Comparison of the current dataset in this study with the only published dataset for the HCC tissue secretome. **B.** Comparison of the enrichment percentage of secretory proteins between our study and that in the human protein database. **C.** Hydrophobicity distribution of the identified secretory proteins. The GO analysis of overlapped different abundance secretory proteins between the comparison of C/DN group and SN/DN group involved biological processes, (**D**) different abundance secretory proteins in the C/DN group involved biological processes, and (**E**) different abundance secretory proteins in the SN/DN group involved biological processes (**F**).

**Figure S2 Representative MS spectra of CA2 in shotgun and targeted proteomics**

**A.** Reporter ion relative intensity of the 8-plex iTRAQ reagents related to CA2 in the MS/MS spectra. **B.** Peak contributions of the individual fragment ions from the unique peptide of CA2. **C.** Representative PRM results of CA2 in HCC patients and healthy volunteers from the training cohort through PRM.

**Figure S3 Representative annotated spectra for the 6 identified peptides of CA2.**

**Figure S4 Validation of the clinical values of serum CA2 concentration**

**A.** The variation tendency of serum CA2 during the follow-up time after radical resection but before recurrence (right), and the sampling time in every patient (left). **B.** TTR and OS were compared between the high- and low-CA2 groups by Kaplan-Meier analysis in the training cohort. **C.** The serum level of CA2 was negatively correlated with the time to recurrence in HCC. **D.** Distribution of serum AFP levels in the validation cohort by ELISA, ***p<0.001. **E.** TTR and OS were compared between the AFP-positive and AFP-negative groups in the validation cohort by Kaplan-Meier analysis. **F.** Distribution of serum AFP levels in AFP-negative HCC patients. **G.** OS was compared between the high and low CA2 groups in AFP-negative HCC patients by Kaplan-Meier analysis.

**Figure S5 Representative images of CA2 immunostaining in tumour tissues**

**Table S1 Basic information and features of HCC patients enrolled in this study.**

**Table S2 The complete list of quantified secretory proteins from this study.**

**Table S3 Differential abundant secretory proteins in C group compared to DN group.**

**Table S4 Differential abundant secretory proteins in SN group compared to DN group.**

**Table S5 Detailed PSM information for the 6 identified peptides of CA2.**

**Table S6 Univariate logistic regression analysis between serum CA2 levels and clinicopathological characteristics.**

## Reference

[1] Ferlay J, Soerjomataram I, Dikshit R, Eser S, Mathers C, Rebelo M, et al. Cancer incidence and mortality worldwide: sources, methods and major patterns in GLOBOCAN 2012. Int J Cancer 2015;136:E359–86.

[2] Wang FS, Fan JG, Zhang Z, Gao B, Wang HY. The global burden of liver disease: the major impact of China. Hepatology 2014;60:2099–108.

[3] Zhou XD, Tang ZY, Yang BH, Lin ZY, Ma ZC, Ye SL, et al. Experience of 1000 patients who underwent hepatectomy for small hepatocellular carcinoma. Cancer 2001;91:1479–86.

[4] Ng KM, Yan TD, Black D, Chu FC, Morris DL. Prognostic determinants for survival after resection/ablation of a large hepatocellular carcinoma. HPB (Oxford) 2009;11:311–20.

[5] Lee YY, McKinney KQ, Ghosh S, Iannitti DA, Martinie JB, Caballes FR, et al. Subcellular tissue proteomics of hepatocellular carcinoma for molecular signature discovery. J Proteome Res 2011;10:5070–83.

[6] Iizuka N, Oka M, Yamada-Okabe H, Nishida M, Maeda Y, Mori N, et al. Oligonucleotide microarray for prediction of early intrahepatic recurrence of hepatocellular carcinoma after curative resection. Lancet 2003;361:923–9.

[7] Sterling RK, Wright EC, Morgan TR, Seeff LB, Hoefs JC, Di Bisceglie AM, et al. Frequency of elevated hepatocellular carcinoma (HCC) biomarkers in patients with advanced hepatitis C. Am J Gastroenterol 2012;107:64–74.

[8] Liu X, Cheng Y, Sheng W, Lu H, Xu Y, Long Z, et al. Clinicopathologic features and prognostic factors in alpha-fetoprotein-producing gastric cancers: analysis of 104 cases. J Surg Oncol 2010;102:249–55.

[9] El-Bahrawy M. Alpha-fetoprotein-producing non-germ cell tumours of the female genital tract. Eur J Cancer 2010;46:1317–22.

[10] Qiao B, Wang J, Xie J, Niu Y, Ye S, Wan Q, et al. Detection and identification of peroxiredoxin 3 as a biomarker in hepatocellular carcinoma by a proteomic approach. Int J Mol Med 2012;29:832–40.

[11] Song IS, Kim HK, Jeong SH, Lee SR, Kim N, Rhee BD, et al. Mitochondrial peroxiredoxin III is a potential target for cancer therapy. Int J Mol Sci 2011;12:7163–85.

[12] Xing X, Huang Y, Wang S, Chi M, Zeng Y, Chen L, et al. Comparative analysis of primary hepatocellular carcinoma with single and multiple lesions by iTRAQ-based quantitative proteomics. J Proteomics 2015;128:262–71.

[13] Feng JT, Liu YK, Song HY, Dai Z, Qin LX, Almofti MR, et al. Heat-shock protein 27: a potential biomarker for hepatocellular carcinoma identified by serum proteome analysis. Proteomics 2005;5:4581–8.

[14] Veenstra TD, Conrads TP, Hood BL, Avellino AM, Ellenbogen RG, Morrison RS. Biomarkers: mining the biofluid proteome. Mol Cell Proteomics 2005;4:409–18.

[15] Anderson NL, Anderson NG. The human plasma proteome: history, character, and diagnostic prospects. Mol Cell Proteomics 2002;1:845–67.

[16] Hanash SM, Pitteri SJ, Faca VM. Mining the plasma proteome for cancer biomarkers. Nature 2008;452:571–9.

[17] Paltridge JL, Belle L, Khew-Goodall Y. The secretome in cancer progression. Biochim Biophys Acta 2013;1834:2233–41.

[18] Slany A, Haudek-Prinz V, Zwickl H, Stattner S, Grasl-Kraupp B, Gerner C. Myofibroblasts are important contributors to human hepatocellular carcinoma: evidence for tumor promotion by proteome profiling. Electrophoresis 2013;34:3315–25.

[19] Yu Y, Pan X, Ding Y, Liu X, Tang H, Shen C, et al. An iTRAQ based quantitative proteomic strategy to explore novel secreted proteins in metastatic hepatocellular carcinoma cell lines. Analyst 2013;138:4505–11.

[20] Cao J, Hu Y, Shen C, Yao J, Wei L, Yang F, et al. Nanozeolite-driven approach for enrichment of secretory proteins in human hepatocellular carcinoma cells. Proteomics 2009;9:4881–8.

[21] Xing X, Liang D, Huang Y, Zeng Y, Han X, Liu X, et al. The application of proteomics in different aspects of hepatocellular carcinoma research. J Proteomics 2016;145:70–80.

[22] Yang L, Rong W, Xiao T, Zhang Y, Xu B, Liu Y, et al. Secretory/releasing proteome-based identification of plasma biomarkers in HBV-associated hepatocellular carcinoma. Sci China Life Sci 2013;56:638–46.

[23] Ma J, Chen T, Wu S, Yang C, Bai M, Shu K, et al. iProX: an integrated proteome resource. Nucleic Acids Res 2019;47:D1211–7.

[24] Chang L, Karin M. Mammalian MAP kinase signalling cascades. Nature 2001;410:37–40.

[25] Gaesser JM, Fyffe-Maricich SL. Intracellular signaling pathway regulation of myelination and remyelination in the CNS. Exp Neurol 2016;283:501–11.

[26] Gherardi E, Birchmeier W, Birchmeier C, Vande Woude G. Targeting MET in cancer: rationale and progress. Nat Rev Cancer 2012;12:89–103.

[27] Veillette A, Grenier K, Brasseur K, Frechette-Frigon G, Leblanc V, Parent S, et al. Regulation of the PI3-K/Akt survival pathway in the rat endometrium. Biol Reprod 2013; 79:1–11.

[28] Solit DB, Basso AD, Olshen AB, Scher HI, Rosen N. Inhibition of heat shock protein 90 function down-regulates Akt kinase and sensitizes tumors to Taxol. Cancer Res 2003;63:2139–44.

[29] Noor SI, Jamali S, Ames S, Langer S, Deitmer JW, Becker HM. A surface proton antenna in carbonic anhydrase II supports lactate transport in cancer cells. Elife 2018;7: e35176.

[30] Chiang WL, Chu SC, Yang SS, Li MC, Lai JC, Yang SF, et al. The aberrant expression of cytosolic carbonic anhydrase and its clinical significance in human non-small cell lung cancer. Cancer Lett 2002;188:199–205.

[31] Mallory JC, Crudden G, Oliva A, Saunders C, Stromberg A, Craven RJ. A novel group of genes regulates susceptibility to antineoplastic drugs in highly tumorigenic breast cancer cells. Mol Pharmacol 2005;68:1747–56.

[32] Haapasalo J, Nordfors K, Jarvela S, Bragge H, Rantala I, Parkkila AK, et al. Carbonic anhydrase II in the endothelium of glial tumors: a potential target for therapy. Neuro Oncol 2007;9:308–13.

[33] Zhou Y, Mokhtari RB, Pan J, Cutz E, Yeger H. Carbonic anhydrase II mediates malignant behavior of pulmonary neuroendocrine tumors. Am J Respir Cell Mol Biol 2015;52:183–92.

[34] Parks SK, Pouyssegur J. Targeting pH regulating proteins for cancer therapy-Progress and limitations. Semin Cancer Biol 2017;43:66–73.

[35] Viikila P, Kivela AJ, Mustonen H, Koskensalo S, Waheed A, Sly WS, et al. Carbonic anhydrase enzymes II, VII, IX and XII in colorectal carcinomas. World J Gastroenterol 2016;22:8168–77.

[36] Zhou R, Huang W, Yao Y, Wang Y, Li Z, Shao B, et al. CA II, a potential biomarker by proteomic analysis, exerts significant inhibitory effect on the growth of colorectal cancer cells. Int J Oncol 2013;43:611–21.

[37] Liu LC, Xu WT, Wu X, Zhao P, Lv YL, Chen L. Overexpression of carbonic anhydrase II and Ki-67 proteins in prognosis of gastrointestinal stromal tumors. World J Gastroenterol 2013;19:2473–80.

[38] Takahashi M, Yang XJ, Sugimura J, Backdahl J, Tretiakova M, Qian CN, et al. Molecular subclassification of kidney tumors and the discovery of new diagnostic markers. Oncogene 2003;22:6810–8.

[39] Hynninen P, Parkkila S, Huhtala H, Pastorekova S, Pastorek J, Waheed A, et al. Carbonic anhydrase isozymes II, IX, and XII in uterine tumors. APMIS 2012;120:117–29.

[40] Liu CM, Lin YM, Yeh KT, Chen MK, Chang JH, Chen CJ, et al. Expression of carbonic anhydrases I/II and the correlation to clinical aspects of oral squamous cell carcinoma analyzed using tissue microarray. J Oral Pathol Med 2012;41:533–9.

[41] Parkkila S, Rajaniemi H, Parkkila AK, Kivela J, Waheed A, Pastorekova S, et al. Carbonic anhydrase inhibitor suppresses invasion of renal cancer cells in vitro. Proc Natl Acad Sci U S A 2000;97:2220–4.

[42] Zhang C, Wang H, Chen Z, Zhuang L, Xu L, Ning Z, et al. Carbonic anhydrase 2 inhibits epithelial-mesenchymal transition and metastasis in hepatocellular carcinoma. Carcinogenesis 2018;39:562–70.

[43] Dunn WB, Broadhurst D, Begley P, Zelena E, Francis-McIntyre S, Anderson N, et al. Procedures for large-scale metabolic profiling of serum and plasma using gas chromatography and liquid chromatography coupled to mass spectrometry. Nat Protoc 2011;6:1060–83.

[44] Liu H, Wang Y, Xing X, Sun Y, Wei D, Chen G, et al. Comparative proteomics of side population cells derived from human hepatocellular carcinoma cell lines with varying metastatic potentials. Oncol Lett 2018;16:335–45.

[45] Song C, Ye M, Han G, Jiang X, Wang F, Yu Z, et al. Reversed-phase-reversed-phase liquid chromatography approach with high orthogonality for multidimensional separation of phosphopeptides. Anal Chem 2010;82:53–6.

[46] Huang X, Zeng Y, Xing X, Zeng J, Gao Y, Cai Z, et al. Quantitative proteomics analysis of early recurrence/metastasis of huge hepatocellular carcinoma following radical resection. Proteome Sci 2014; 22:1–14.

