## Supplementary figures and images for "Quantitative Analysis of Tissue Secretome Reveals the Diagnostic and Prognostic Value of Carbonic Anhydrase II in Hepatocellular Carcinoma"

### Supplementary Figure 1

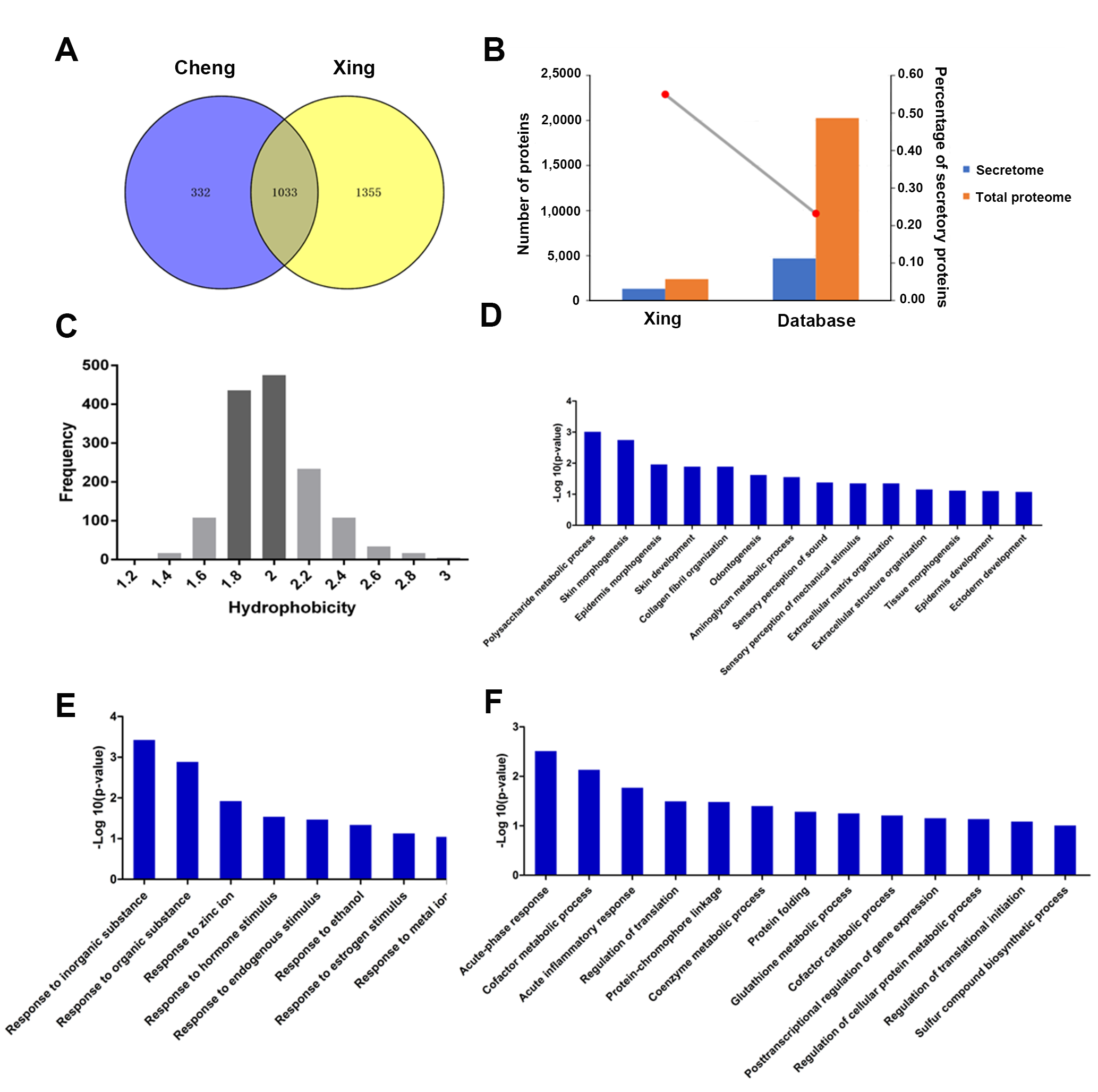

### Supplementary Figure 2

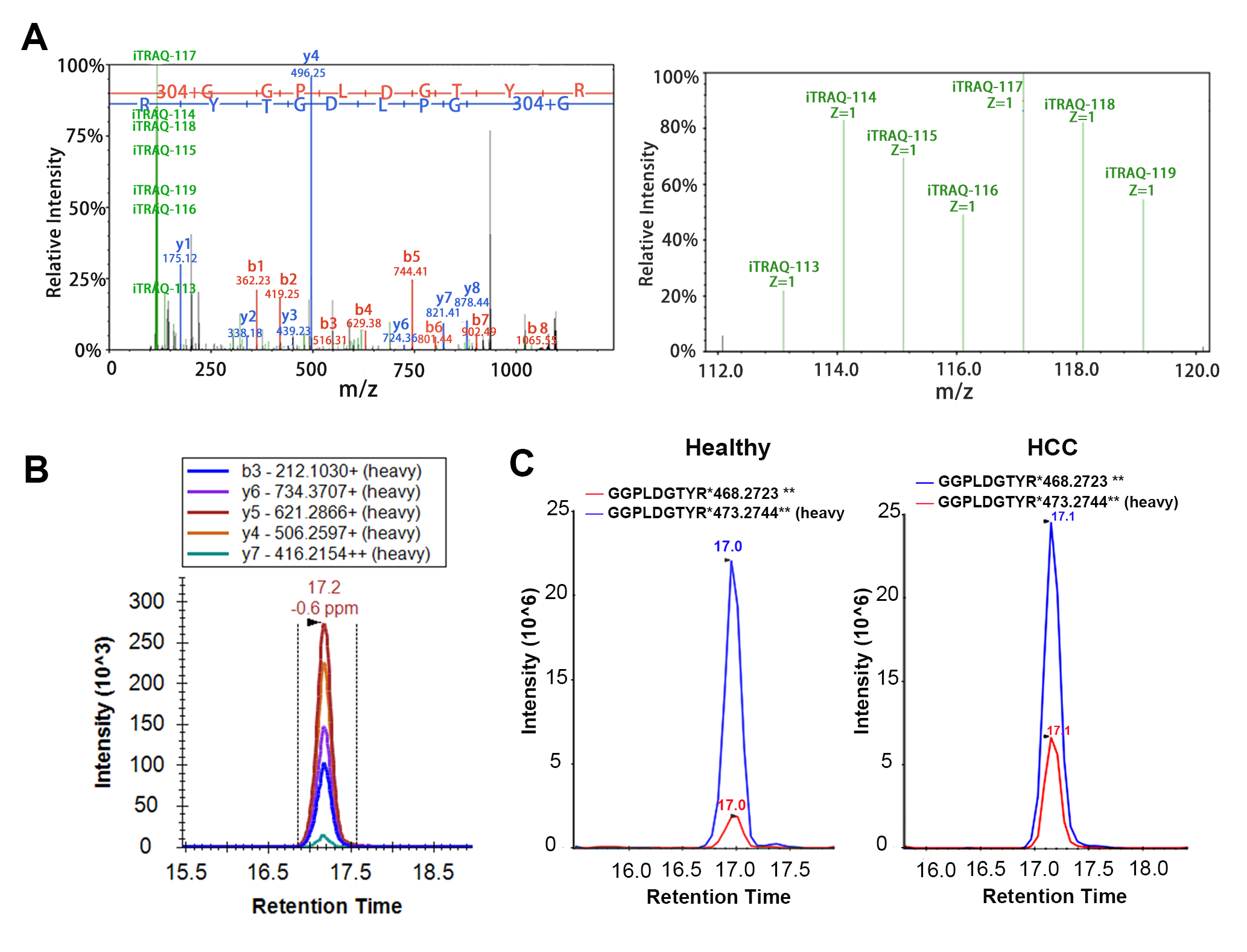

### Supplementary Figure 3

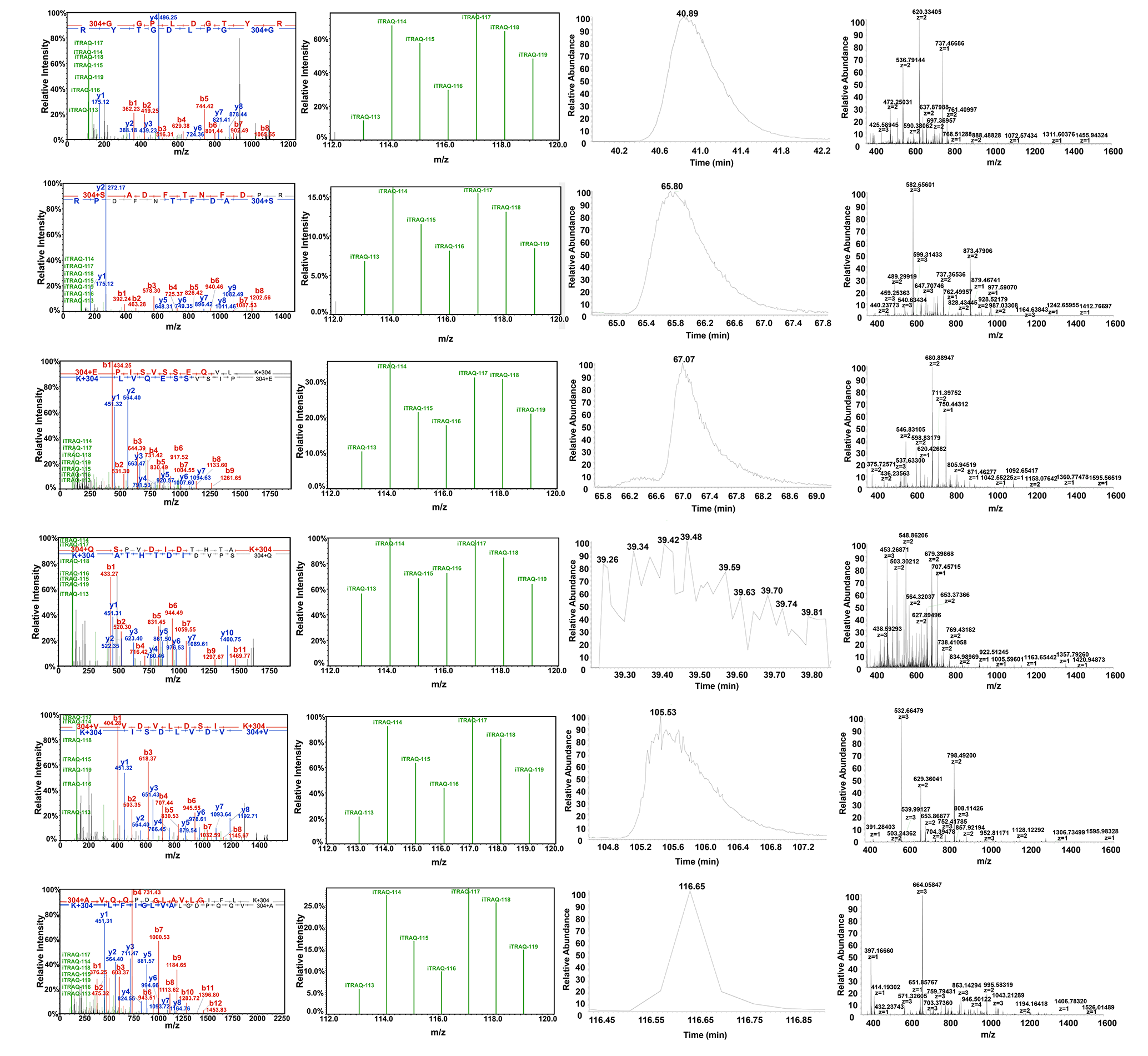

### Supplementary Figure 4

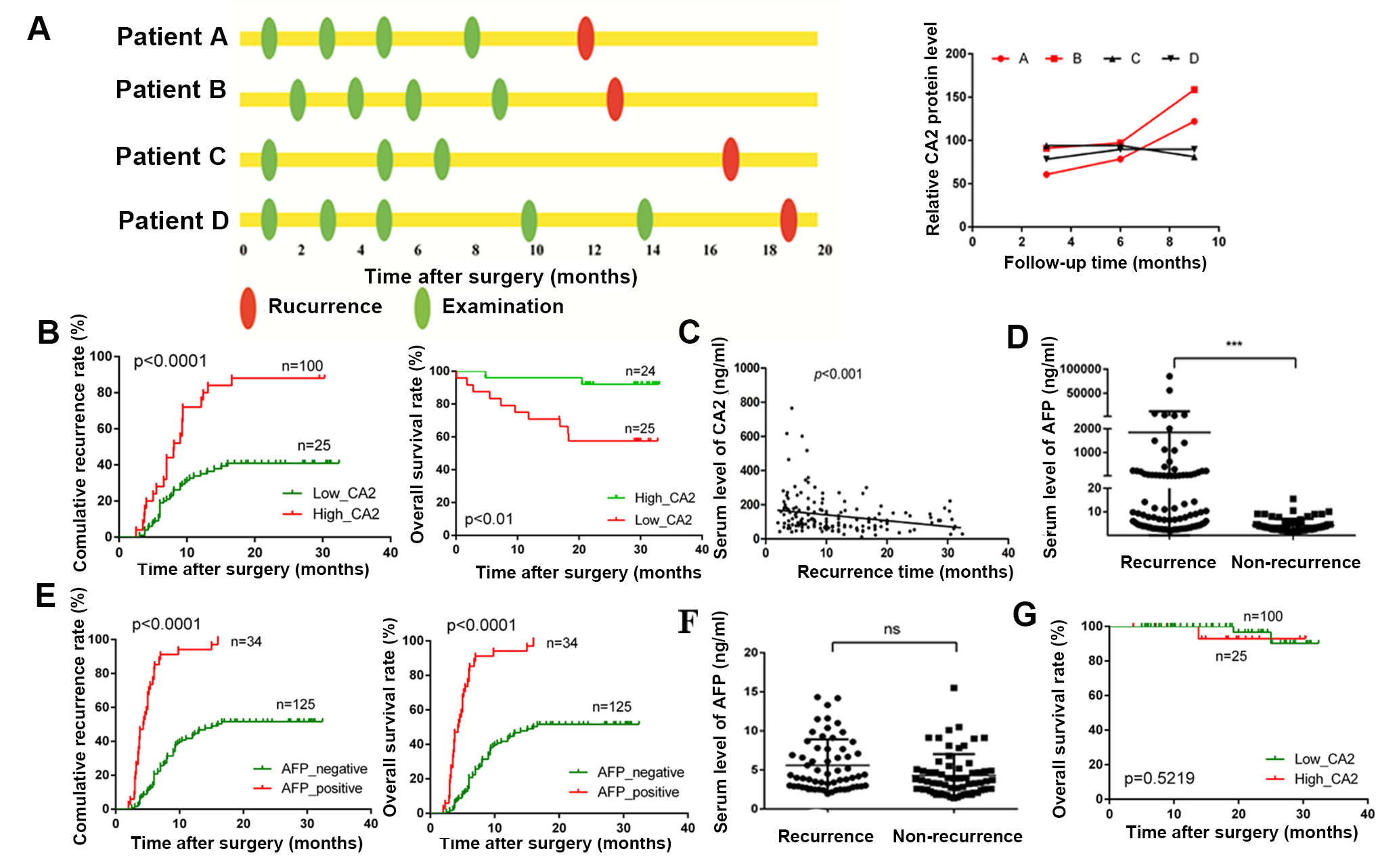

### Supplementary Figure 5

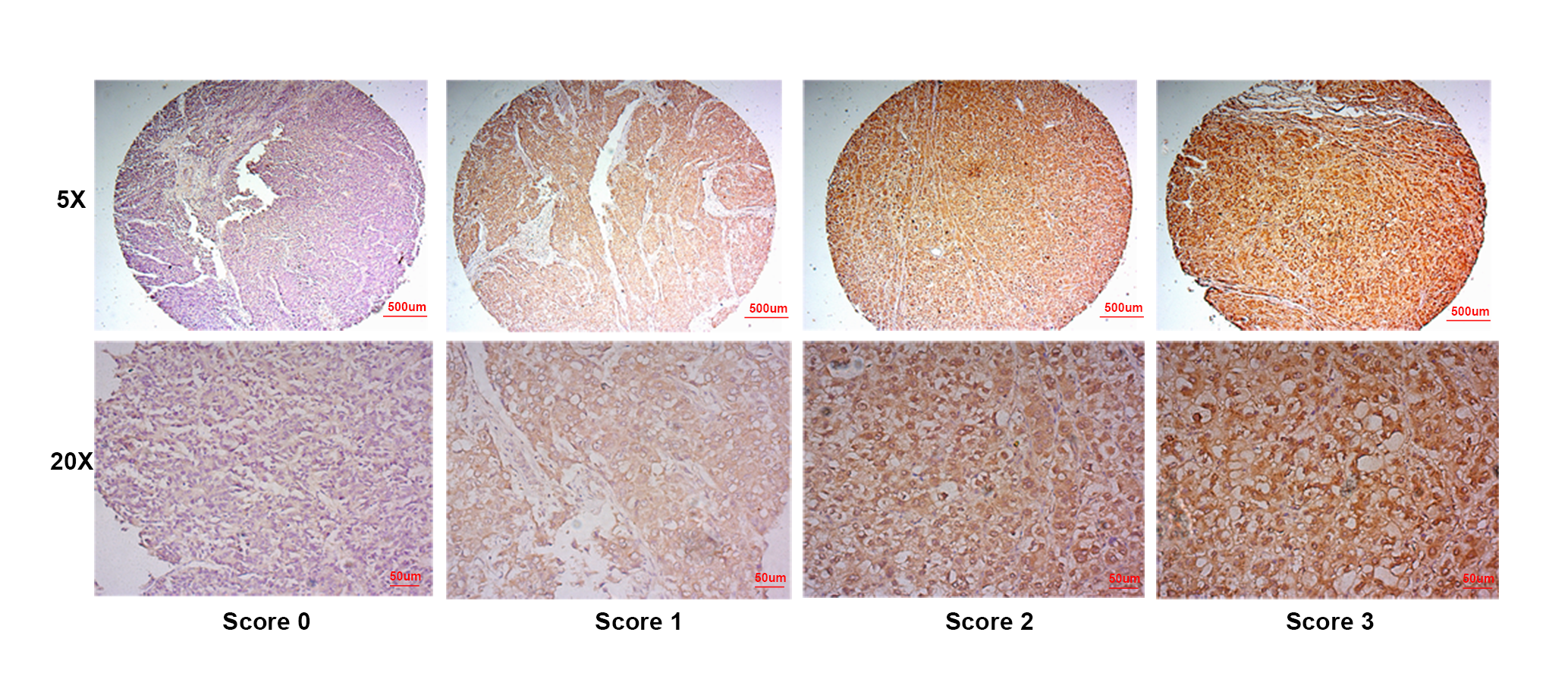
